# Genetic diversity and population structure of *Phytophthora infestans* in Guatemala and Honduras

**DOI:** 10.1101/2025.05.28.656702

**Authors:** Myriam Izarra, Willmer Perez, Emil Vasquez, Glenda Perez, Soledad Gamboa, Jorge Andrade-Piedra, Breny Flores, Luz Montejo, Arie Sanders, Jan Kreuze

## Abstract

*Phytophthora infestans,* the causal agent of late blight, remains a major constraint to potato production worldwide. In Central America, particularly in Guatemala and Honduras, information on the pathogen’s population structure is limited despite its economic importance and reliance on imported seed potatoes. This study characterized the population structure of *P. infestans* from potato fields in Guatemala and Honduras using microsatellite markers, mitochondrial haplotypes, and mating type analyses. Four SSR-defined groups were identified, corresponding to the clonal lineages EU_13_A2, US_25, US_8, and a distinct group designated CeA (Central America). EU_13_A2 was predominant, accounting for 141 of the 154 isolates analyzed, whereas US_25, CeA, and US_8 were represented by eight, four, and one isolate, respectively. Reference isolates from Nicaragua previously designated as NI_1 clustered with US_25, while reference isolates from Honduras designated as HN_1 clustered with EU_13_A2. A total of 89 multilocus genotypes were identified, indicating substantial SSR genotypic diversity despite the predominance of EU_13_A2. All successfully characterized isolates exhibited the Ia mitochondrial haplotype. Hierarchical AMOVA showed that most SSR variation occurred within departments (69.86%, *p=*0.002), whereas variation among departments within countries (15.97%, *p*=0.092) and between Honduras and Guatemala (14.17%, *p =*0.108) was not statistically significant. The predominance of EU_13_A2 together with the occurrence of additional genetically distinct groups highlights the importance of continued monitoring of *P. infestans* populations and their movements across Central America.

## Introduction

*Phytophthora infestan*s (Mont.) de Bary is an oomycete responsible for late blight in potatoes (*Solanum tuberosum* L.), tomatoes (*Solanum lycopersicum* L.), and other Solanaceae species (Perez & Forbes, 2010). The disease is a well-known primary biotic limitation affecting potato cultivation on a global scale and has been the cause of devastating epidemics, such as the one leading to the Irish Potato Famine of 1845, and still poses a significant challenge to global food security (Day & Shattock, 1997; Dong & Zhou, 2022). Management of the disease with fungicides continues to be the most common approach (Abuley & Hansen, 2022), with a limited number of alternatives available to chemical control (Kang & Dobinson, 2004).

*P. infestans* is a heterothallic species that requires mating types A1 and A2 for sexual reproduction, resulting in oospore formation (Drenth et al., 1995). This pathogen combines the advantages of sexual cycles, which generate genetic diversity, and an expansive population through asexual reproductive cycles (McDonald & Linde, 2002). In sexual populations, both sexual and asexual propagules act as inoculum sources, whereas asexual populations rely entirely on inoculum generated through asexual reproduction (Flier et al., 2002). Monitoring the A1/A2 ratio is crucial to understand the potential for sexual recombination, which supports their evolution (Barton & Charlesworth, 1998). Sexual reproduction allows *P. infestans* to persist in the soil between seasons as thick-walled oospores (Andersson, 2007). In addition, *P. infestans* strains derived from sexual recombination could potentially exhibit a greater capacity to adapt rapidly to environmental fluctuations, including fungicide selection pressure (Goodwin, 1997). The emergence and global spread of aggressive, metalaxyl-resistant clonal lineages such as EU13A2 further illustrate this adaptive potential (Cooke et al., 2012; Islam et al., 2023). Furthermore, in *P. infestans,* genetic diversity is fostered by adaptive evolution, which is driven by the pathogen’s unique genomic structure, particularly in regions enriched with rapidly evolving effector genes (Haas et al., 2009; Leesutthiphonchai et al., 2018; Zhang et al., 2019; Coomber et al., 2024).

The diversity of *P. infestans* is examined by using a combination of phenotypic and genotypic markers. Microsatellite markers, also referred to as simple sequence repeats (SSRs), are frequently used to determine multilocus genotypes (MLGs) and evaluate genetic diversity within populations (Li et al., 2012; Martin et al., 2019). SSR markers are highly polymorphic, affordable, reproducible, neutral, and co-dominant (Lees et al., 2006), and can differentiate between genotypes, even among clonal lineages (Hansen et al., 2016). Mitochondrial haplotypes are used to trace maternal lineages and provide insights into population structure and migration patterns (Griffith & Shaw, 1998; Widmark et al., 2007; Guo J et al., 2009).

Mexico has been proposed as the geographic center of origin of late blight based on several lines of evidence, including the historical occurrence of both mating types (A1 and A2) and the remarkabl e diversity of *P. infestans* in central Mexico in terms of virulence characteristics and molecular markers (Gallegly, 1958; Fry et al., 1993). However, an alternate theory proposes Peru (or the Andean region) as the center of origin, based on phylogenetic, genomic, and population genetic evidence, including the global prevalence of the US-1 clonal lineage, which is notably absent in Mexico (Abad & Abad, 1997; Gomez-Alpizar et al., 2007; Saville et al., 2016; Martin et al., 2016; Coomber et al., 2024). A recent study conducted by Patarroyo *et al*. (2024), based on approximate Bayesian computation of alternative migration scenarios, identified a Peruvian origin of *P. infestans* as the most probable scenario among those evaluated, with an initial migration toward Colombia and Mexico, followed by subsequent migration from Mexico to the United States and later to Europe and Asia, with no return to northern South America.

The A1 mating type dispersed worldwide during the Irish potato famine, whereas the A2 mating type was first reported in Europe, the Middle East, and South America in 1981 (Spielman et al., 1991; Goodwin, 1995a). Genomic studies of historical herbarium samples have shown that the lineage responsible for the nineteenth-century famine pandemic, HERB-1, was genetically distinct from current populations (Yoshida et al., 2013; Saville et al., 2016). The related FAM-1 lineage was widely distributed across multiple continents during the nineteenth and early twentieth centuries before being displaced by later clonal lineages, including US-1, which became globally dominant during the mid-twentieth century (Saville & Ristaino, 2021). In the United States, US-1 was subsequently displaced by newly emerged A2 clonal lineages, including US_8, during the early 1990s, increasing the risk of late blight epidemics (Goodwin, 1995b; Goodwin et al., 1998). More recently, the A2 clonal lineage US_25 has been reported only from New York State (Ristaino, 2026). In Central America, the NI_1 clonal lineage, characterized by the A2 mating type and Ia mitochondrial haplotype, was reported as a predominant lineage infecting both potato and tomato in Nicaragua (Blandón-Díaz et al., 2012).

Guatemala produces over 540,000 metric tons of potatoes per year, which are primarily exported to the neighboring El Salvador, and less than 1% of the crop is planted with certified seed potatoes. Late blight is the most important disease because of its cool and wet weather conditions (Joyce, 2020), and there is high genetic variability among isolates (Ruiz-Chutan et al., 2018). The United States is a significant supplier of seed potatoes to Guatemala, and few quantities have been imported from Peru (World Bank, 2023a; b). In contrast, potato cultivation in Honduras was concentrated in Intibucá, Ocotepeque, La Paz, and Francisco Morazán, where 39,633 metric tons of potatoes were produced between 2011 and 2012. The Honduran potato industry also relies on imports; in 2014 and 2015 6,000 tons were used annually for industrial processing, while seed potato imports reached 1,000 tons. Similar to Guatemala, late blight is recognized as the most significant disease affecting potato cultivation in Honduras (Toledo, 2016). According to the USDA, in 2021, the value of potato imports to Honduras grew by 1.9%, increasing from US$7.8 million in 2016 to US$8.4 million in 2020. Most potatoes for fresh consumption were imported from the United States, whereas most seed potatoes were imported from the Netherlands, France (SAG & UPEG, 2021; World Bank, 2023c), Guatemala, and Chile (Sanders et al., 2024).

This research focused on examining the population structure of *Phytophthora infestans* that impacts potato crops in Guatemala and Honduras. We specifically investigated the composition of clonal lineages, the distribution of mating types, mitochondrial haplotypes, and multilocus genetic diversity to create a regional baseline for epidemiological monitoring and informed disease management strategies.

## Materials and Methods

### Sampling and sample processing

The sampling strategy used in this study was designed to identify dominant pathogen genotypes in Honduras and Guatemala. Between 2021-2022, potato leaflet samples exhibiting late blight symptoms were collected from the main potato-growing areas at altitudes ranging from 1170 to 2088 m above sea level (masl) in Honduras and 2140.57–3699 masl in Guatemala, with approximately 53 fields studied based on latitude and longitude (Table S1).

Samples were collected on FTA cards (Whatman FTA Classic Card, catalog number WB120055; GE Healthcare UK Ltd.), following the manufacturer’s instructions (Cooke, 2020a). Each sample had its sporulating side facing down, crushed individually in the field, and then FTA cards were stored in zip-lock bags and sent by post for processing in the pathology laboratory at the International Potato Center (CIP), Lima, Peru.

DNA extraction from FTA was performed following the manufacturer’s instructions, with some modifications using two discs of 3 mm, and two rinses with modified Tris(hydroxymethyl) aminomethane-ethylenediaminetetraacetic acid (TE^−1^) buffer (10 mM Tris and 0.1 mM EDTA), and then resuspended in 50 µL TE buffer (10mM Tris and 1mM EDTA) and incubated for 5 min at 95 °C as the last step before being stored at −20 °C until use.

### Mating type determination

The mating types were determined using the W16 CAPS assay to distinguish A1 and A2 mating types of *P. infestans* isolates using PCR primers W16-1 (5’-AACACGCACAGGCATATAAATGTA-3’) and W16-2 (5’-GCGTAATGTAGCGTAACAGCTCTC-3’) following the protocol outlined by Judelson et al. (1995). The PCR reaction was carried out in 20 μL mixture, which included 4 μL of 5x buffer (Gotaq®, Promega), 6 μl MgCl_2_ (25 mM), 0.2 μl dNTPs (100 μM), 0.4 μM of each primer, 1 U of Taq DNA polymerase (Gotaq®, Promega) and 10ng of template DNA. The thermal cycling conditions were as follows: initial denaturation at 94 °C for 5 min, followed by 30 cycles of denaturation at 94 °C for 1 min, primer annealing at 53 °C for 1 min, and primer elongation at 72 °C for 1 min. The PCR process was concluded with a final elongation step at 72 °C for 10 minutes. Subsequently, the PCR products were subjected to a three-hour digestion at 37 °C using the restriction enzyme (recognition site 5ü-GG/CC-3ü from New England Biolabs (NEB) (https://www.neb.com). The digestion mixture for each sample included 7.7 μL of H_2_O, 2 μL of CutSmart Buffer 10X, 0.3 μL *Hae III* (NEB, #R0108) and 10 μl of PCR product (Mazáková et al., 2006; Brylińska et al., 2018). Mating type determination was based primarily on the presence or absence of the undigested 557 bp fragment following *Hae III* digestion. The mating type A2 was confirmed using the PHYB-1 (5’-GAT CGG ATT AGT CAG ACG AG-3ü) and PHYB-2 (5’-GCG TCT GCA AGG CGC ATT TT-3ü) primers (Kim & Lee). The PCR was performed in 20 μL mixture, which included 4 μL of 5x buffer (Gotaq®, Promega), 1.2 μL MgCl_2_ (25 mM), 0.2 μl dNTPs (2.5 mM), 0.4 μM of each primer, 1 U of Taq DNA polymerase (Gotaq®, Promega) and 10ng of template DNA. The thermal cycling conditions were as follows: initial denaturation at 95 °C for 5 min, followed by 35 cycles of denaturation at 94 °C for 1 min, primer annealing at 62 °C for 1 min, and primer elongation at 72 °C for 1 min. The PCR process was concluded with a final elongation step at 72 °C for 7 minutes. Mating type assignments were based on concordant results obtained from both the W16 CAPS and the A2-specific PHYB marker.

### Mitochondrial haplotype determination

Mitochondrial DNA (mtDNA) haplotypes were determined following the profiles of Griffith and Shaw (1998). This process included amplifying the P2 and P3 mitochondrial regions which were then digested with *MspI* for P2 and *EcoR*I for P3. The combined restriction DNA patterns from P2/*MspI* and P3/*EcoRI* enabled differentiation among haplotypes Ia, Ib, II a, and IIb. The resulting restriction fragment length polymorphisms (RFLPs) were separated through gel electrophoresis and compared to reference profiles - for precise haplotype classification.

### Microsatellite analysis

The genomic DNA of *the P. infestans* isolates was amplified using a predetermined set of microsatellite markers following a modified version of the protocol for the 12-plex method (Table S1, Li et al., 2013). The Type-It Microsatellite PCR kit (QIAGEN, https://www.qiagen.com) was used for PCR reactions according to a modified protocol described by Saville *et al*. (2016). The PCR amplicons were then sent to Arizona State University for post-PCR processing in 96-well plates. To prepare the samples for fragment analysis on the 3730 instrument, 3 μL of the LIZ500 size standard ladder was mixed with 1 mL of formamide in a 96-sample plate. Each well of the plate contained 9 μL of the ladder/formamide mixture and 2 μL of 1:50 diluted sample. The total sample volume in each well was 11 μL when loaded onto 3730 instrument for fragment analysis.

### SSR allele scoring and genotype structure

Microsatellite genotypes were obtained using a 12-plex SSR system for Phytophthora infestans, following standard protocols (Cooke, et al., 2012; Li et al., 2013). SSR allele sizing was conducted using GeneMarker v.1.9 software (SoftGenetics, State College, PA, USA), applying allele binning criteria described by Li et al. (2013). Previously characterized reference isolates representing EU_13_A2, US_8 and US_25, obtained from The James Hutton Institute (Dundee, UK), TBAS (Coomber et al., 2023) and the Fry laboratory (Cornell University), were included to standardize allele scoring and facilitate lineage assignment. Allele calls were manually reviewed following the EuroBlight SSR allele-scoring guidelines (Cooke, 2020b), considering locus-specific stutter patterns, relative peak intensity, amplification quality, and minor size-calling variation.

For each isolate, up to three observed alleles per locus were retained. Loci with fewer than three observed alleles were retained as recorded, without imputation of additional alleles or allele dosage. Unobserved allele positions were coded as 0 in the SSR dataset and treated as missing information in downstream analyses. Multilocus genotypes (MLGs) were defined from the observed allele profiles across the 12 SSR loci using mlg.id() in the *poppr* package (Kamvar et al., 2014).

Pairwise genetic distances among isolates were calculated using Bruvo’s genetic distance based on multilocus SSR profiles. Bruvo’s distance accounts for the stepwise mutation model of microsatellite evolution and allows comparison of genotypes with varying ploidy (Bruvo et al., 2004).

Clonal lineage assignments were based on comparison of field-isolate SSR profiles with reference isolates representing EU_13_A2, US_25, and US_8. Lineage interpretation considered multilocus SSR similarity and genetic relationships observed in the PCoA and Neighbor-Joining analyses based on Bruvo’s genetic distance. Additional previously characterized reference genotypes from the Americas and Europe were included in the Neighbor-Joining analysis to provide a broader regional context for lineage comparison. Field isolates clustering consistently with a reference lineage were assigned to the corresponding lineage.

### Data Analysis

Statistical analyses were performed using R version 4.6.1 (R Core Team, 2026). The *poppr* package was used to calculate multilocus genotypes (MLGs), diversity statistics, and other population genetic parameters (Kamvar et al., 2014). Locus-level diversity statistics, including Simpson’s diversity (1-D), expected heterozygosity (Hexp), and evenness, were calculated using locus_table() after removing zero-coded unobserved allele positions using recode_polyploids().

Population grouping varied according to the objective of each analysis. For the PCoA, isolates were grouped according to their sampling locations, while reference isolates representing known clonal lineages were retained for lineage comparison. For the hierarchical analysis of molecular variance (AMOVA), populations were defined geographically, with departments nested within countries. Genetic variation was partitioned among countries, and within departments. Clone correction was applied within departments before AMOVA, retaining one representative of each MLG, and statistical significance was evaluated using 999 permutations.

Reference isolates were excluded from population genetic diversity and the MLG-based minimum spanning network but were retained in the PCoA and Neighbor-Joining analyses to facilitate comparison with previously characterized clonal lineages.

Population genetic relationships were evaluated by principal coordinates analysis (PCoA) based on Bruvo’s genetic distance (Bruvo et al., 2004) using the *polysat* package (Clark & Jasieniuk, 2011). A Neighbor-Joining tree was reconstructed using the BioNJ algorithm from a matrix of Bruvo’s genetic distances calculated for field isolates and reference genotypes (Saitou & Nei, 1987; Gascuel, 1997). In addition to the EU_13_A2, US_25, and US_8 reference isolates included in the PCoA, additional previously characterized reference genotypes from the Americas and Europe were included to provide a broader geographic context for lineage comparison. The resulting tree was visualized and annotated using the Interactive Tree of Life (iTOL)(Letunic & Bork, 2024).

Genotypic relationships were further explored using a minimum spanning network (MSN), constructed with the poppr package (Kamvar et al., 2014), based on Bruvo’s genetic distance. Each MLG was represented as a node, with node size reflecting the number of isolates sharing that MLG. Edge width and shading represent Bruvo’s genetic distance between connected MLGs.

Genotypic diversity based on MLG frequencies was evaluated according to sampling location. Simpson’s and Shannon–Wiener diversity indices were used to characterize MLG diversity, while genotypic richness was estimated as the expected number of multilocus genotypes (eMLG), standardized to the smallest sample size using rarefaction (Simpson, 1949; Shannon, 2001).

## Results

### Samples collected

In Honduras, 144 potato samples were collected from three departments (equivalent to the state): Intibucá, Francisco Morazán and Ocotepeque. In Guatemala, 10 potato samples were collected from four departments: Huehuetenango, San Marcos, Chimaltenango, and Sololá (Tables 1 and S2).

### Mating type test

The primer set W16–1 and W16–2 generated a 557-bp product. Following digestion with *HaeIII*, A2 isolates were identified by the absence of the undigested 557-bp fragment and the presence of the expected 457-bp fragment; the complementary 100-bp fragment was observed in several representative samples (Fig. S1A). Using this assay, the 154 isolates showed the molecular pattern associated with A2.

For confirmation, amplification with the PHYB-1/PHYB-2 primers produced a 347-bp fragment in isolates from Honduras (n=143) and Guatemala (n=9), supporting their assignment to the A2 mating type (Fig. S1B). A total of 152 isolates were concordantly identified as A2 by both marker systems (Table S2). Because only isolates with concordant results between the W16 CASPS assay and the A2-specific PHYB marker were considered confidently assigned, all isolates meeting these criteria were molecularly assigned to the A2 mating type; the remaining 2 isolates could not be conclusively assigned because the PHYB region was not successfully amplified.

### Genotypic characterization

Mitochondrial haplotype analysis identified 144 isolates as haplotype Ia (Table S2). Digestion of the P2 region with *MspI* produced fragments of 720 and 350 bp, a restriction pattern consistent with the haplotype Ia or IIb (Fig. S2A). To distinguish between these haplotypes, the P3 region was digested with *EcoRI*. Successful amplification of the P3 region was obtained for 153 isolates (143 from Honduras and 10 from Guatemala), yielding fragments of 1064 and 228 bp, corresponding to group I haplotypes (Fig. S2B). The combined P2/*MspI* and P3/*EcoRI* restriction patterns identified all 166 successfully analyzed isolates as haplotype Ia; the remaining 1 isolate could not be conclusively assigned because the P3 region was not successfully amplified.

Allelic diversity varied among the 12 SSR loci (Table 2). The number of alleles per locus ranged from 2 to 17, with a mean of 5.42. D13 was the most polymorphic locus, with 17 alleles and the highest values of Simpson’s diversity (1-D = 0.83) and expected heterozygosity (Hexp=0.83). In contrast, Pi70 showed the lowest diversity (1-D=0.01; Hexp=0.01). Across loci, mean Simpson’s diversity, expected heterozygosity, and evenness were 0.45, 0.46, and 0.71, respectively (Table 2).

**Table 1.** Number of *Phytophthora infestans* isolates collected in Honduras and Guatemala.

| **Country** | **Departments** | **Number of isolates** | **Fields** | **Altitud range (masl)** |
|---|---|---|---|---|
| **Honduras** | Intibucá | 137 | 41 | 1170-2088 |
|  | Ocotepeque | 4 | 1 | 1900 |
|  | Francisco Morazán | 3 | 1 | 1558 |
| **Guatemala** | San Marcos | 2 | 2 | 2836.9-2886.5 |
|  | Chimaltenango | 1 | 1 | 2140.55 |
|  | Huehuetenango | 6 | 6 | 3009.6-3699 |
|  | Sololá | 1 | 1 | 2447.2 |
| **Grand Total** |  | 154 | 53 |  |

**Table 2.** Genetic diversity statistics for the 12 microsatellite loci analyzed in *Phytophthora infestans* populations from Honduras and Guatemala.

| **locus** | **allele** | **1-D** | **Hexp** | **Evenness** |
|---|---|---|---|---|
| D13 | 17 | 0.83 | 0.83 | 0.65 |
| SSR8 | 5 | 0.59 | 0.59 | 0.84 |
| SSR4 | 9 | 0.60 | 0.60 | 0.71 |
| Pi04 | 3 | 0.51 | 0.51 | 0.94 |
| Pi70 | 2 | 0.01 | 0.01 | 0.33 |
| SSR6 | 4 | 0.54 | 0.54 | 0.86 |
| Pi63 | 3 | 0.54 | 0.54 | 0.89 |
| G11 | 7 | 0.64 | 0.64 | 0.73 |
| SSR3 | 4 | 0.44 | 0.45 | 0.80 |
| SSR11 | 4 | 0.17 | 0.17 | 0.43 |
| SSR2 | 2 | 0.06 | 0.06 | 0.44 |
| Pi4B | 5 | 0.52 | 0.52 | 0.89 |
| mean | 5.42 | 0.45 | 0.46 | 0.71 |

The hierarchical AMOVA based on geographic structure indicated that most of the genetic variation occurred within departments (69.86%, p=0.002). Variation among departments within countries accounted for 15.97% of the total genetic variation but was not statistically significant (p=0.092). Similarly, differences between Honduras and Guatemala accounted for 14.17% of the total genetic variation and were not statistically significant (p=0.108) (Table 3).

**Table 3.** Hierarchical analysis of molecular variance (AMOVA) based on SSR data, with departments nested within countries, for *Phytophthora infestans* populations from Honduras and Guatemala.

|  |  |  |  | Components of covariance |  | Monte-Carlo test for significance nrepet=999 |  |  |  |
|---|---|---|---|---|---|---|---|---|---|
|  | Df | SumSq | MeanSq | Sigma | % | Obs | Std.Obs | Alter | Pvalue |
| Within departments | 87 | 57.05 | 0.66 | 0.66 | 69.86 | 0.66 | -4.87 | less | 0.002 |
| Among departments within countries | 5 | 6.11 | 1.22 | 0.15 | 15.97 | 0.15 | 1.37 | greater | 0.092 |
| Between countries | 1 | 4.73 | 4.73 | 0.13 | 14.17 | 0.13 | 1.40 | greater | 0.108 |
| Total | 93 | 67.89 | 0.73 | 0.94 | 100 |  |  |  |  |

Genetic relationships among isolates based on microsatellite data and Bruvo genetic distance were explored using PCoA. The analysis revealed four main groups, with most isolates from Honduras (n=137) and Guatemala (n=4) clustering with the EU13A2 reference isolates. A second group was associated with the US_25 reference isolates, and a third group with US_8 reference isolates, and a fourth distinct group lacked a corresponding reference genotype (Figure 1).

**Figure 1.**
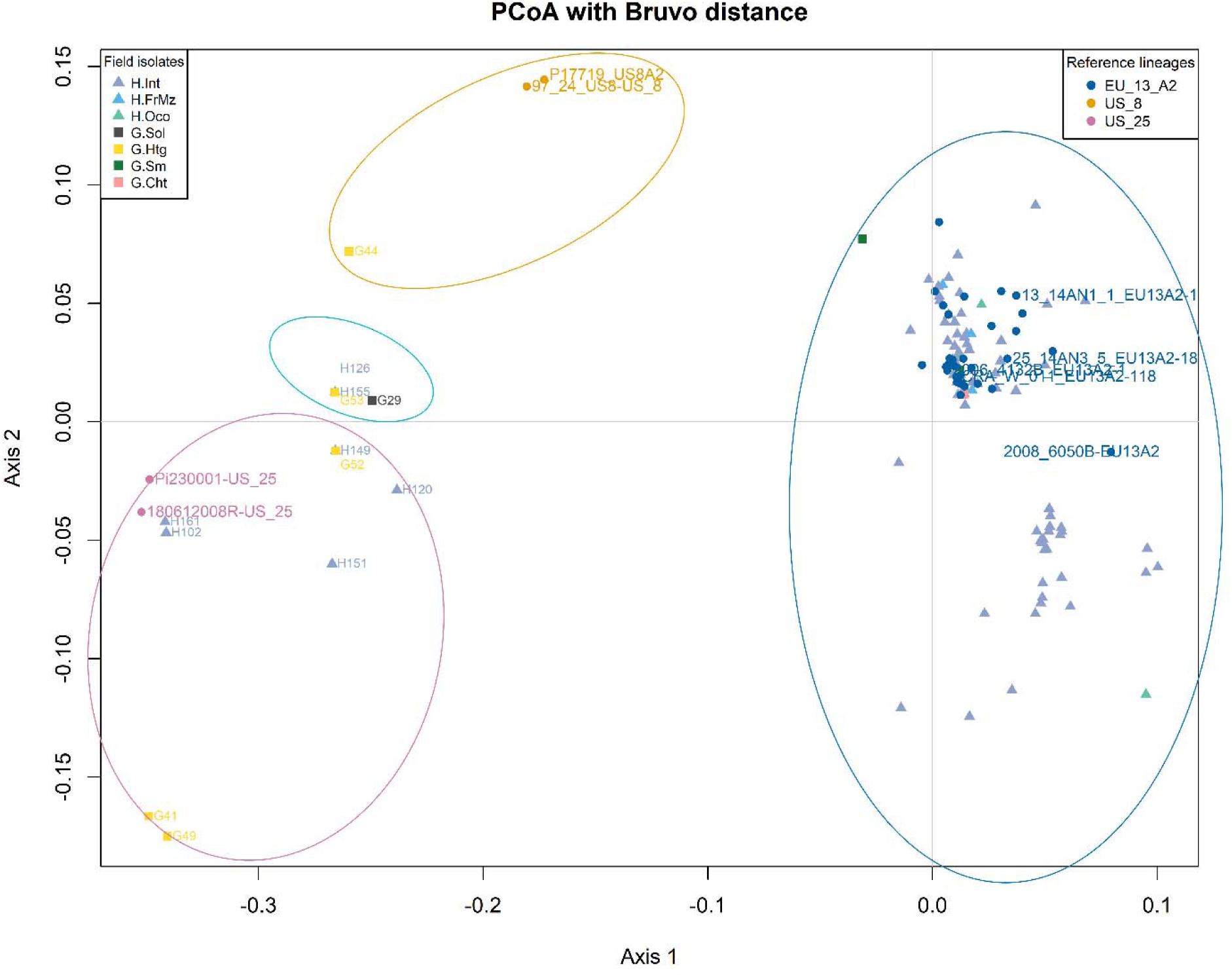
Principal coordinates analysis (PCoA) based on Bruvo’s genetic distance of *Phytophthora infestans* isolates from Honduras and Guatemala. The prefix H denotes Honduras and G denotes Guatemala. Triangles represent isolates from Honduras and squares represent isolates from Guatemala. Isolates are colored according to their sampling location: H.Int (Intibucá, light lilac), H.FrMz (Francisco Morazán, sky blue), H.Oco (Ocotepeque, light greenG.Sol (Sololá, dark gray), G.Qtz (Quetzaltenango, dark red), G.Htg (Huehuetenango, yellow), G.Sm (San Marcos, dark green), and G.Cht (Chimaltenango, pink). Reference isolates representing clonal lineages EU_13_A2 (dark blue), US_8 (orange), and US_25 (magenta) are indicated. Ellipses highlight genetic groups among isolates.

The Neighbor-Joining tree based on Bruvo’s genetic distance separated the field isolates into four main clusters associated with the identical clonal lineages (Figure 2). The cluster associated with EU_13_A2 comprised 141 isolates and was predominantly represented by isolates from Honduras, including Intibucá (n=130), Ocotepeque (n=4), and Francisco Morazán (n=3), together with isolates from Guatemala collected in San Marcos (n=2), Chimaltenango (n=1), and Huehuetenango (n=1). Previously characterized reference isolates from Honduras designated as HN_1 clustered within the EU_13_A2 group together with EU_13_A2 reference isolates from Europe. Reference isolates from Nicaragua previously designated as NI_1 clustered within the US_25 group together with the US_25 reference isolate. The CeA cluster comprised four field isolates from Intibucá (n=2), Huehuetenango (n=1), and Sololá (n=1), whereas US_8 was represented by a single field isolate from Huehuetenango, Guatemala. Additional reference genotypes from Americas and Europe provided broader geographic context for comparison among the identified clonal lineage (Figure 2). The geographic distribution of the four lineages is shown in Figure 3.

**Figure 2.**
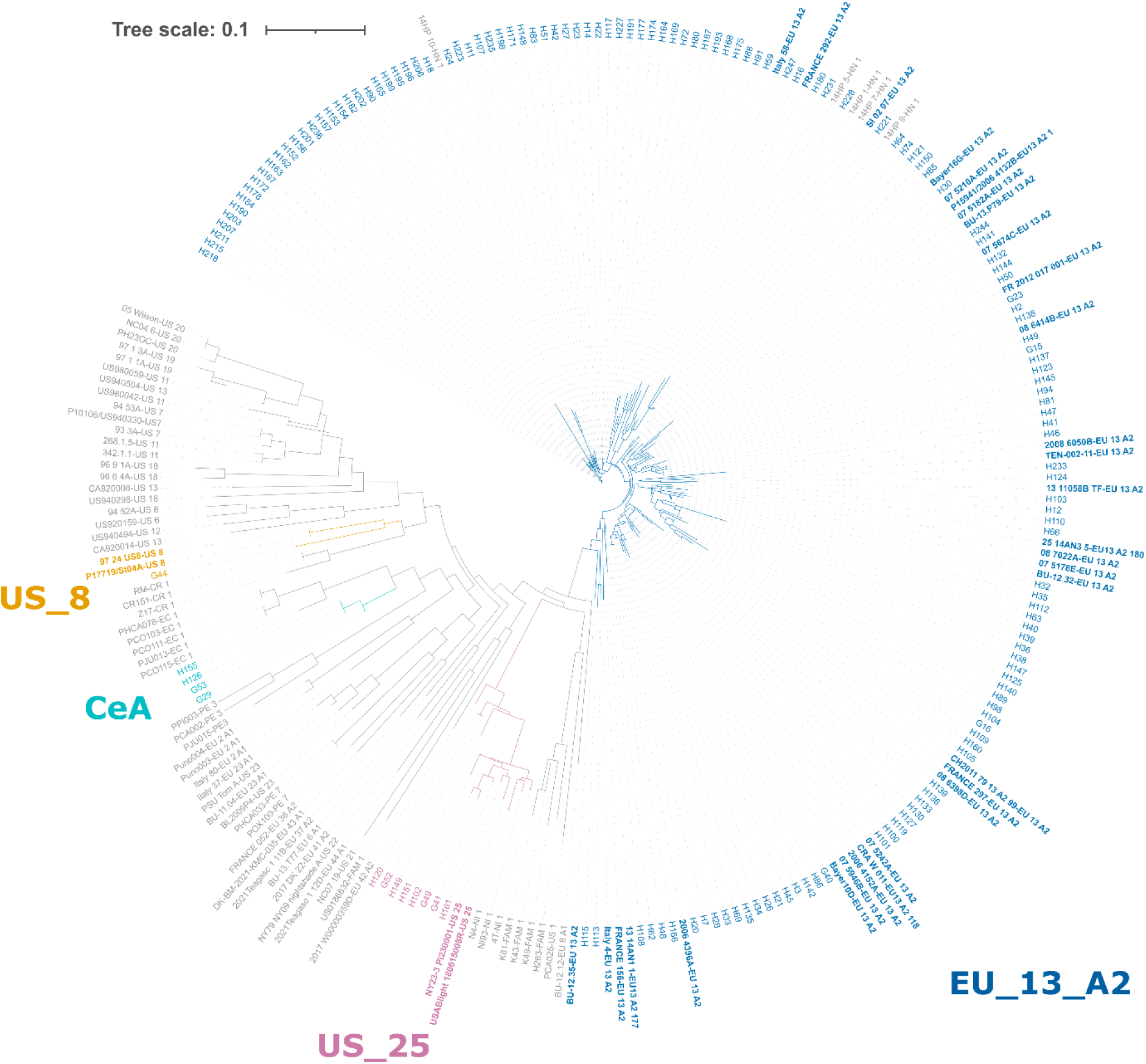
Neighbor-Joining (BioNJ) tree based on Bruvo genetic distance showing the relationship among *Phytophthora infestans* isolates from Honduras and Guatemala and reference genotypes. Reference isolates representing previously characterized clonal lineages from Europe and the America were included for comparison and lineage assignment. The EU_13_A2, US_25 and US_8 clonal lineage are shown in dark blue, magenta, and orange, respectively, while the CeA cluster is shown in cyan.

**Figure 3.**
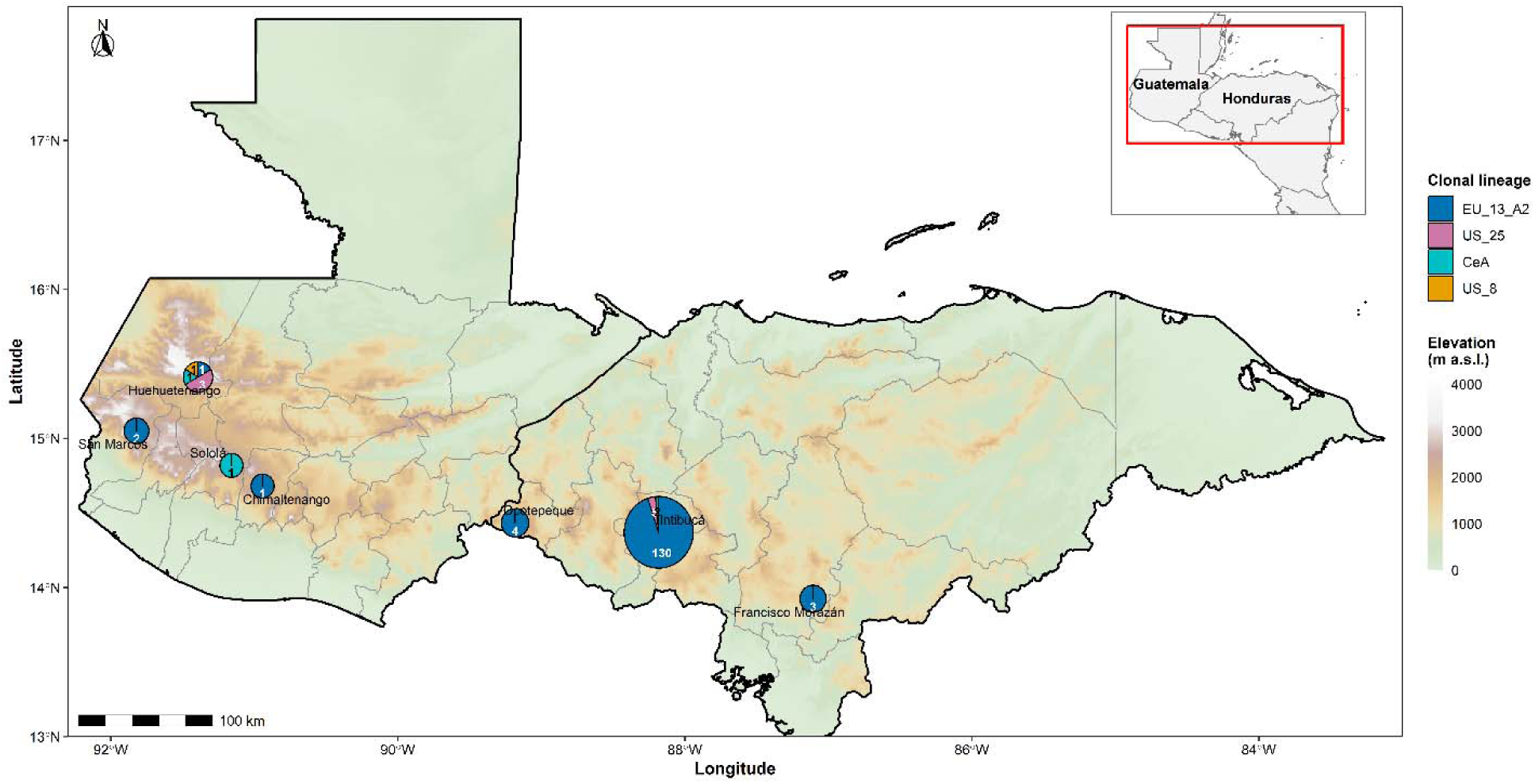
Geographical distribution of *Phytophthora infestans* clonal lineages EU_13_A2, US_25, CeA, and US_8 among isolates (n =154) collected in Honduras and Guatemala in 2022. Pie charts represent the proportion of isolates assigned to each clonal lineage per department, with pie-chart size proportional to the total number of isolates sampled. Number within pie-chart sectors indicate the number of isolates assigned to each clonal lineage. Background shading represents elevation in meters above sea level (m.a.s.l.), and departmental boundaries are shown in gray. The international boundary line between Guatemala and Honduras (right) is indicated by a black line. The inset shows the location of Guatemala and Honduras within Central America and highlights the geographic extent the study area.

A total of 89 multilocus genotypes (MLGs) were identified among the 154 isolates analyzed. The allele profiles of the 89 MLGs across the 12 SSR loci are provided in Table S3. Diversity analyses excluded reference isolates. Intibucá, which represented the largest sampled population (n=137), contained 77 MLGs whereas the remaining sampling locations had considerably smaller sample sizes, ranging from one to six isolates (Table 4).

Across the complete dataset, genotypic diversity was high (H=4.13; λ=0.97), with an evenness value (E.5) of 0.62 and an expected heterozygosity (Hexp) of 0.46 (Table 4).

Significant multilocus linkage disequilibrium was detected in the pooled dataset (IA = 2.923, p=0.001; rbarD = 0.313, p=0.001), indicating non-random association among loci.

Of the 89 MLGs identified, 68 were represented by a single isolate. The most frequent genotypes were MLG53 (n=14), MLG68 (n=10), and MLG60 (n=8). EU_13A2 comprised 141 isolates representing 79 MLGs, whereas US_25 comprised eight isolates representing seven MLGs. The four CeA isolates represented two MLGs, while the single US_8 isolate represented a unique MLG. Four MLGs were shared between Honduras and Guatemala: MLG75 and MLG80 within EU_13_A2, MLG7 within US_25, and MLG86 within CeA. The minimum spanning network (MSN) illustrated genetic relationships among the 89 MLGs, with closely related MLGs occurring within the assigned clonal lineages (Figure S3).

**Table 4.** Population genetic diversity statistics based on 12 microsatellite (SSR) loci for *Phytophthora infestans* isolates grouped according to sampling location in Honduras and Guatemala. *H:Honduras, G:Guatemala, Int: Intibucá, Oco: Ocotepeque, FrMz: Francisco Morazán, Sm: San Marcos, Cht: Chimaltenango, Sol: Sololá, Htg: Huehuetenango.

| **Pop** | **N** | **MLG** | **eMLG** | **SE** | **H** | **G** | **lambda** | **E.5** | **Hexp** | **Ia** |
|---|---|---|---|---|---|---|---|---|---|---|
| H.Int | 137 | 77 | 9.11 | 0.89 | 3.98 | 33.5 | 0.97 | 0.62 | 0.44 | 2.31 |
| H.Oco | 4 | 4 | 4 | 0 | 1.39 | 4 | 0.75 | 1 | 0.47 | 0.59 |
| H.FrMz | 3 | 3 | 3 | 0 | 1.10 | 3 | 0.67 | 1 | 0.54 | 1 |
| G.Sm | 2 | 2 | 2 | 0 | 0.69 | 2 | 0.5 | 1 | 0.58 | NA |
| G.Cht | 1 | 1 | 1 | 0 | 0 | 1 | 0 | NaN | 0.75 | NA |
| G.Sol | 1 | 1 | 1 | 0 | 0 | 1 | 0 | NaN | 0.75 | NA |
| G.Htg | 6 | 6 | 6 | 0 | 1.79 | 6 | 0.83 | 1 | 0.59 | 1.80 |
| Total | 154 | 89 | 9.23 | 0.84 | 4.13 | 38.6 | 0.97 | 0.62 | 0.46 | 2.92 |

*N* number of individuals, *MLG* number of multilocus genotypes, *eMLG* expected number of MLGs, *SE* Standard error. The subsequent columns are descriptors of MLG diversity: *H* Shannon- Weiner Index, *H Stoddart and Taylor's Index (G), lambda* Simpson's index, *E.5* Evenness, *H<sub>exp</sub>* Nei's unbiased gene diversity. *IA* Index of Association, *rbarD* standardized index of association.

## Discussion

The primary finding of this study was the predominance of the EU13A2 clonal lineage in the *Phytophthora infestans* population sampled in Honduras and Guatemala, accounting for 141 of the 154 field isolates analyzed. EU13A2 is a widespread and epidemiologically important lineage that has caused severe late blight outbreaks in Europe and Asia (Cooke et al., 2012; Chowdappa et al., 2013). Assignment of the Central American isolates to EU_13_A2 was supported by their genetic relationships with multiple EU_13_A2 reference isolates included in the SSR analyses. Notably, historical reference isolates from Honduras designated as HN_1, collected in 2010 and 2014 (Coomber et al., 2023) clustered within the EU_13_A2 group together with European EU_13_A2 references. This close relationship suggests that EU_13_A2-related genotypes were already present in Honduras before the present sampling, although the available SSR data cannot establish when or through which route this lineage was introduced into Central America.

Despite the predominance of EU_13_A2, considerable genotypic variation was detected within this lineage, with 79 MLGs identified among 141 EU_13_A2 isolates. Variation among SSR profiles within established clonal lineages of *P. infestans* has been reported previously and may accumulate through mutation and other processes during clonal propagation (Goodwin et al., 1994; Stellingwerf et al., 2018; Göre et al., 2021). Intralineage SSR variation has also been documented specifically within 13_A2 populations. In South India, minor variants of 13_A2 were distinguished by variation at the highly variable D13 and SSR4 loci (Chowdappa et al., 2015), consistent with subclonal variation previously reported for 13_A2 in Europe (Cooke et al., 2012). Notably, D13 was also the most polymorphic locus in the present study, with 17 alleles. The relatively high SSR variation observed within EU_13_A2 in Central America may therefore reflect intralineage diversification. However, the present data cannot determine whether the observed variation arose after introduction into Central America or was already present in the introduced population. Multiple introductions of closely related genotypes may also have contributed to the observed diversity. Genome-wide markers would provide greater resolution for determining the relationships among these EU_13_A2 genotypes and their relationship to populations from other geographic regions.

Three additional genetic groups, US_25, CeA and US_8, were detected at substantially lower frequencies. US_25 comprised eight field isolates from Honduras and Guatemala. This lineage was first detected in New York state in 2018 and has subsequently been reported only from limited areas of New York (Ristaino, 2026). Notably, reference isolates from Nicaragua previously designated as NI_1 clustered with the US_25 reference isolate and with the Central American field isolates assigned to this group. This close similarity based on the 12-locus SSR profiles suggests that US_25-related genotypes may have a broader geographic distribution than previously recognized. However, higher-resolution genomic analyses would be required to determine the evolutionary relationship between the Nicaraguan NI_1 isolates, the US_25 genotype reported in the United States, and the isolates identified in Honduras and Guatemala. CeA comprised four isolates and remained differentiated from the reference genotypes included in the SSR analyses. Although CeA shares some SSR allele sizes with EC-1, it did not cluster with the EC-1 references and carried mitochondrial haplotype Ia rather than the I Ia haplotype reported for EC-1 (Martin et al., 2016; Lindqvist-Kreuze et al., 2020), supporting its recognition here as a distinct putative genetic group. US_8 was represented by only one isolate from Guatemala. Genotypes related to US_8 have previously been reported in Central America, including Costa Rica (Gomez-Alpizar, 2005; Lucca, Florencia et al., 2023), although the occurrence of genetically similar SSR profiles does not by itself demonstrate direct epidemiological connections among these populations.

All isolates confidently characterized for mating type were assigned to A2, contrasting with earlier reports from Honduras in which A1 isolates were detected (Forbes, 2004). Changes in mating-type composition have accompanied major shifts in *P. infestans* populations in other regions following the introduction and expansion of new lineages (Drenth et al., 1993; Deahl et al., 2008). The predominance of A2 in the present samples may therefore reflect temporal changes in population composition and the expansion of A2 lineages, particularly EU_13_A2, although differences in sampling intensity and the possible persistence of A1 at low frequency cannot be excluded. Moreover, the occurrence of A2 should not itself be interpreted as evidence of sexual reproduction. Because conventional pairing assays with known A1 and A2 tester isolates were not performed, the potential occurrence of self-fertile isolates also cannot be evaluated solely from the molecular assays used here.

The mitochondrial results showed considerably less variation than the nuclear SSR markers. All successfully characterized isolates carried mitochondrial haplotype Ia, consistent with previous reports from the region, including the replacement of haplotype IIb by Ia in Nicaraguan potato fields near the Honduran border (Blandón-Díaz et al., 2012) and with reference haplotypes characterized by Martin *et al*. (2019). The occurrence of a single mitochondrial haplotype across genetically differentiated SSR groups demonstrates that the mitochondrial uniformity does not necessarily correspond to nuclear genetic uniformity. Consequently, the predominance of haplotype Ia should not be interpreted as evidence for a single introduction into the region, because distinct lineages or genotypes may share the same mitochondrial background.

The geographic AMOVA provided little evidence for strong differentiation between the sampled regions. Most SSR variation occurred within departments (69.86%, *p*=0.002), whereas variation among departments within countries accounted for 15.97% of the total variation and was not statistically significant (*p*= 0.092). Similarly, variation between Honduras and Guatemala accounted for 14.17% and was not statistically significant (p=0.108). These results suggest that genetic variation is primarily distributed at the local level rather than being strongly partitioned according to national or departmental boundaries. The predominance of EU_13_A2 in the sampled population is consistent with this limited geographic differentiation. Furthermore, four identical MLGs were detected in both Honduras and Guatemala (MLG7, MLG75, MLG80, and MLG 86), representing EU_13_A2, US_25 and CeA, and providing additional evidence that some multilocus genotypes are shared across national boundaries. Movement of infected planting material and other propagules could contribute to the distribution of related genotypes across geographic boundaries, but the present SSR data do not provide sufficient resolution to reconstruct specific introduction or dispersal routes between Honduras and Guatemala. Therefore, whether EU_13_A2 entered the two countries independently or spread regionally following one or more introductions remains unresolved.

The widespread occurrence of EU_13_A2 in the present study has important implications for late blight management in Central America. Movement of infected seed potatoes has been recognized as an important pathway for long-distance dissemination of *P. infestans*, and the distribution of clonal lineages in several regions has been associated with movement of planting material (Montarry et al., 2010; Beninal et al., 2022). EU_13_A2 has also demonstrated the capacity to expand rapidly after introduction into new geographic regions, as reported in India, where it became an important component of late blight populations (Dey et al., 2018). Honduras relies substantially on imported seed potatoes, including material originating from European countries where EU_13_A2 was first reported (Cooke et al., 2012; Mariette et al., 2015; SAG & UPEG, 2021; World Bank, 2023c). Imported seed potatoes therefore represent a plausible historical pathway for the introduction of EU_13_A2-related genotypes, although the present study cannot establish the source or direction of introduction. The predominance of EU_13_A2 may have important implications for late blight management because this lineage has previously been characterized as highly aggressive, capable of overcoming resistance in some potato cultivars (Lees et al., 2012; Cooke et al., 2012). Metalaxyl-resistant populations of 13_A2 have also been reported in South India (Chowdappa et al., 2013, 2015).

The coexistence of EU_13_A2 with US_25, CeA and US_8 further highlights the potential for movement and persistence of genetically differentiated *P. infestans* groups in Central America. Infected seed tubers can provide primary inoculum early in the growing season and facilitate disease establishment and subsequent spread (Johnson, 2010). Strengthening phytosanitary surveillance of imported planting material, monitoring changes in lineage composition, and supporting the production and distribution of high-quality seed potatoes could therefore contribute to reducing the risk of introduction and dissemination of epidemiologically important genotypes. Strategies to strengthen local seed systems may include early-generation seed production, decentralized multiplication, and appropriate quality-assurance mechanisms (Forbes et al., 2020).

Overall, our results demonstrate a *P. infestans* population in Honduras and Guatemala strongly dominated by EU_13_A2 but containing substantial intralineage SSR variation and several additional genetic groups. The association of historical HN_1 reference genotypes for interpreting historical lineage designations across Central America. The absence of significant differentiation between countries, together with the predominance of EU_13_A2 across the sampled area, highlights the need for coordinated regional surveillance rather than monitoring populations solely within national boundaries. Future studies incorporating genome-wide-markers, broader geographic sampling, and phenotypic characterization of aggressiveness, virulence, and fungicide sensitivity will be important for resolving introduction histories and determining the epidemiological significance of the genetic variation observed here.

## Author contributions

MI: coordination, sample processing, molecular laboratory work, marker scoring, analysis, interpretation of data, and drafting of the manuscript.

WP: coordination.

SG: sample processing.

JA: coordination.

EV: sampling.

BF: sampling.

AS: coordination.

GP: sampling.

LM: sampling.

JK: interpretation of data and drafting of the manuscript.

## Supplementary Information

### Supplementary Table Legend

**Table S1.** Details of the SSR markers of the 12-plex microsatellite assay utilized for the characterization of *Phytophthora infestans* isolates

**Table S2.** Phenotypic, genotypic, geographic, and sampling information for *Phytophthora infestans* isolates from Honduras and Guatemala, including multilocus genotype (MLG) assignments and allele profiles across the 12 SSR loci

**Table S3.** Multilocus genotypes (MLGs) identified among *Phytophthora infestans* isolates from Honduras and Guatemala and their allele profiles across 12 SSR loci.

### Supplementary Figure Legend

**Figure S1.** Mating type compatibility assay of Honduran and Guatemalan *Phytophthora infestans* isolates. (**A**) *HaeIII*-digested W16 amplicons; (**B**) PCR products generated with PHYB primers from Honduran (H7, H120) and Guatemalan (G53, G44) isolates. 1 kb Plus DNA ladder.

**Figure S2.** RFLP-based mitochondrial haplotypes of Honduran and Guatemalan *Phytophthora infestans* isolates. (**A**) *MspI* digestion of the P2 region; (**B**) *EcoRI* digestion of the P3 region from Honduran (H7, H120) and Guatemalan (G53, G44) isolates. 1 kb Plus DNA ladder.

**Figure S3.** Minimum spanning network (MSN) of the 89 multilocus genotypes (MLGs) identified among 154 *Phytophthora infestans* field isolates from Honduras and Guatemala, based on Bruvo’s genetic distance. Each node represents an MLG, with node size proportional to the number of isolates sharing that MLG. Node colors indicate the assigned clonal lineages (EU_13_A2, US_25, CeA, and US_8). Edge width and shading represent Bruvo’s genetic distance between connected MLGs.

## Acknowledgements

We would like to thank Freddy Ventura for technical support during sample processing. We also thank Dr. David Cooke, Dr. Ristaino, M.S Saville and Dr. Hansen for providing the raw data used as controls for the scoring process. Field collections in Honduras and Guatemala were conducted in collaboration with the DICTA and ICTA, respectively. We would like to thank both institutes for their excellent support and contributions.

## Funder Information Declared

This study was funded by the CGIAR Plant Health Initiative (PHI), funded by the CGIAR Trust Fund Donors (https://www.cgiar.org/funders/) and the Feed the Future Innovation Lab for Current and Emerging Threats to Crops provided by the United States Agency for International Development (USAID) cooperative agreement No: 7200AA21LE00005

## Data Availability Statement

The generated data are available on the Supplementary Table S2, available at https://www.biorxiv.org/content/10.1101/2025.05.28.656702v4

## Footnotes

Conflict of Interest declaration: The authors declare that they have no affiliations with or involvement in any organization or entity with any financial interest in the subject matter or materials discussed in this manuscript.

