## Supplementary material for "Genetic diversity and population structure of *Phytophthora infestans* in Guatemala and Honduras": Figure S1, Figure S2, Figure S3

**Supplementary Information**

**Supplementary Table Legend**

**Table S1**. Details of the SSR markers of the 12-plex microsatellite assay utilized for the characterization of *Phytophthora infestans* isolates

**Table S2**. All data of the *Phytophthora infestans* isolates characterized in this study

**Supplementary Figure**

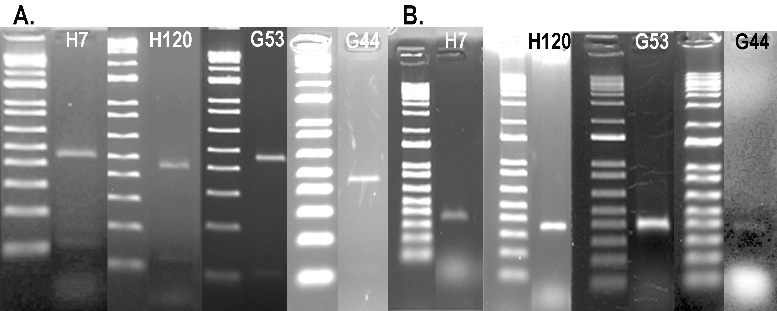

**Figure S1**. Mating type compatibility assay of Honduran and Guatemalan *Phytophthora infestans* isolates. (**A**) HaeIII-digested W16 amplicons; (**B**) PCR products generated with PHYB primers from Honduran (H7, H120) and Guatemalan (G53, G44) isolates. 1 kb Plus DNA ladder.

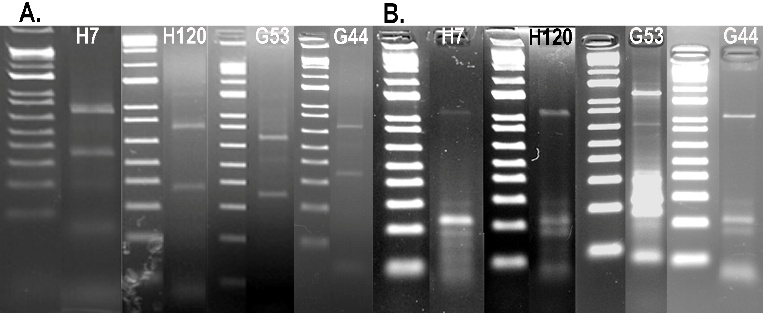

**Figure S2**. RFLP-based mitochondrial haplotypes of Honduran and Guatemalan *Phytophthora infestans* isolates. **(A)** MspI digestion of the P2 region; **(B)** EcoRI digestion of the P3 region from Honduran (H7, H120) and Guatemalan (G53, G44) isolates. 1 kb Plus DNA ladder.

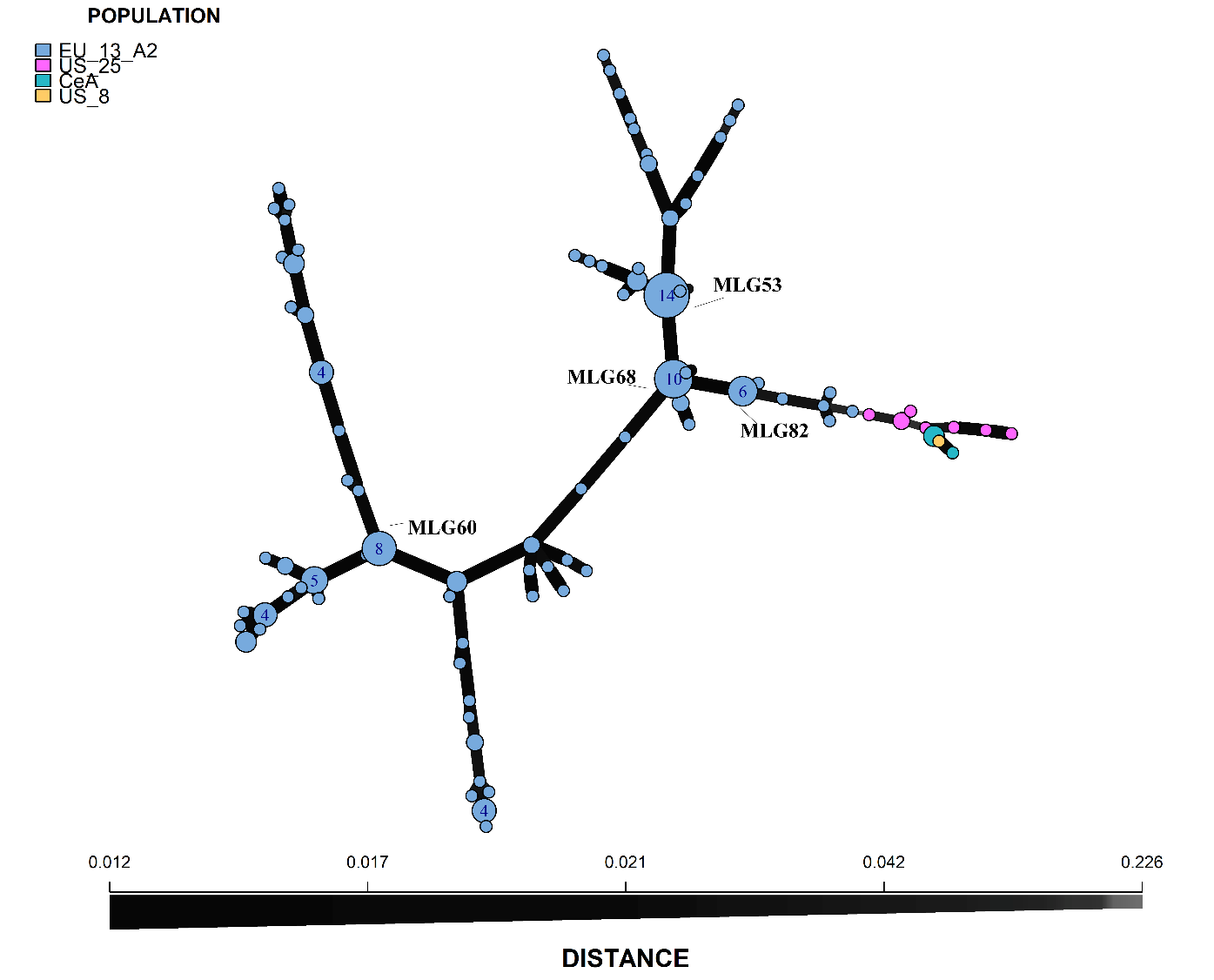

**Figure S3**. Minimum spanning network (MSN) of the 89 multilocus genotypes (MLGs) identified among 154 *Phytophthora infestans* field isolates from Honduras and Guatemala, based on Bruvo’s genetic distance. Each node represents an MLG, with node size proportional to the number of isolates sharing that MLG. Node colors indicate the assigned clonal lineages (EU_13_A2, US_25, CeA, and US_8). Edge width and shading represent Bruvo’s genetic distance between connected MLGs.
